# The Role of Edible Habitat Complexity in Food Webs

**DOI:** 10.64898/2026.03.23.712465

**Authors:** Eden J. Forbes, Jason D. Stockwell

## Abstract

Habitat complexity (HC) in part determines the diversity, stability, and behavior of food webs and can influence predation according to a wide variety of functional relationships. Many aquatic species provide habitat complexity and are also consumed by other species (e.g., macrophytes, corals, mussels). However, food web theory does not readily account for these species that act as ‘edible habitat complexity’ (EHC). Here, we combine existing theory on predator-prey interactions, HC, and prey switching to describe the role of EHC in benthic food web models. We dissect feedback loops in each model to demonstrate how self-regulation of the prey species, mediated by species densities and HC, drives that food web’s behavior. HC can stabilize predator-prey interactions by coupling prey self-regulation with HC self-regulation. EHC can further stabilize predator-prey interactions across a wide variety of “HC functions” that relate HC to predation rates.

**Significance Statement:** Habitat complexity (HC) plays a critical role in trophic interactions, population dynamics, and food web stability. However, little theory exists to describe ‘edible habitat complexity’ (EHC), where a species is both consumed and confers habitat complexity for other species. We provide a series of models demonstrating how HC and EHC alter the population dynamics and stability of simple aquatic food webs. HC is strongly stabilizing in food webs by providing ‘safety in rarity’ for prey. EHC provides safety in rarity for both prey and the EHC species because their predators are omnivorous. Given the prevalence of EHC species in aquatic systems (e.g., macrophytes, corals, mussels), our models demonstrate the importance of maintaining EHC species in aquatic systems for stable food webs.

## INTRODUCTION

Habitat complexity (HC) impacts food webs across organizational levels via trophic interactions and their functional responses (Crowder and Cooper 1982, Diehl 1992, Soukup et al. 2022). Most frequently, HC has been connected to biodiversity (Simpson 1949, Tews et al. 2003) and predation risk (Savino and Stein 1982, Boukal et al. 2007), either increasing or inhibiting the ability of predators to capture prey. HC’s influence on predation rates (henceforth, “HC-predation function”) ranges from linear relationships to binary presence/absence effects (Mocq et al. 2021). Intermediate non-linear HC-predation functions are common (Tokeshi and Arakaki 2012) and demonstrate the ‘threshold hypothesis’, where predation rates rapidly change at some HC density (Gotceitas and Colgan 1989). Still, the role of HC in food web theory is less clear (Kovalenko et al. 2012). We do know HC can reduce over-exploitation of prey by predators by stabilizing predator-prey dynamics (Scheffer and De Boer 1995, van Nes and Scheffer 2005) and non-linear HC-predation functions can support alternative ecosystem states (Scheffer et al. 1993). Still, less is known about the mechanisms underlying stability changes and how outcomes change with different HC-predation functions.

Moreover, many sources of HC are themselves edible (‘edible habitat complexity’, EHC) such as macrophytes and corals. While the contributions of EHC species to community composition are documented (Crowder and Cooper 1982, Thomaz et al. 2008, Christie et al. 2009, Komyakova et al. 2013), EHC is rarely discussed as its own biotic compartment and its impact on food webs is unclear (but see Lee 2006, Filbee-Dexter and Sheibling 2014). For example, Dreissenid mussels represent another prevalent form of EHC. Zebra (*Dreissena polymorpha*) and quagga (*D. bugensis*) mussels impact benthic communities in three significant ways. First, they create HC, lowering predation of mid-trophic predators on benthic prey (Beekey et al. 2004, McCabe et al. 2006). Second, dreissenids enrich benthic food webs by filtering biomass from the water column, increasing production of benthic invertebrates 3-to 5-fold (Stewart et al. 1998a, b). Third, dreissenids themselves are prey, most notably for round goby *Neogobious melanostromus* (Lederer et al. 2006, Lederer et al. 2008). Gobies do not prefer dreissenids (Diggins et al. 2002, Foley et al. 2017) but can significantly reduce dreissenid populations in the absence of preferred prey (Barton et al. 2005, Naddafi and Rudstam 2014). Understanding how round goby predation affects *Dreissena* abundance is an ongoing need (Ruetz et al. 2012) as no general theory predicts the consequences of round goby invasions (Meerhoff and Gonzalez-Sagrario, 2022). Describing the principles of EHC in food webs is an important step.

We demonstrate the potential role of EHC using three benthic food web models. The first is the canonical consumer-resource model (Rosenzweig 1971) with logistic growth of invertebrates and type II predation by a mid-trophic predator. The second model includes *Dreissena* in two life history stages, juveniles and mature adults. Mature dreissenids act as HC, reducing predation risk of benthic invertebrates. The predator does not consume *Dreissena* in this second model (e.g., sculpins, *Cottidae*). The third model renders *Dreissena* edible (EHC) with round goby as the predator. Specifically, juvenile *Dreissena* are edible because round gobies do not consume mature dreissenids (Ghedotti et al. 1995, Andraso et al. 2011). Goby predators thus switch between consuming invertebrates and juvenile dreissenids, while mature dreissenids provide refuge for invertebrates.

We used these models to explore how HC and EHC affect benthic food webs. As parameterized, HC couples invertebrate self-regulation to juvenile *Dreissena* self-regulation, limiting conditions under which invertebrate self-regulation creates positive feedback. To demonstrate, we employed ‘loop tracing’, the dissection of feedback loops that underlie food web stability (e.g. Dambacher et al. 2003, Novak et al. 2016, Cortez 2024, Forbes and Hall 2025). We also show that the specific HC-predation function, varying in inflection point and steepness, can stabilize or destabilize food webs. Then, we use loop tracing to show that EHC stabilizes food webs across a broad range of HC-predation functions.

## MODEL DESCRIPTION

### Predator-Prey Model

A standard consumer-resource model governs the trophic interaction between benthic invertebrates (*I*) and their predators (*P*):

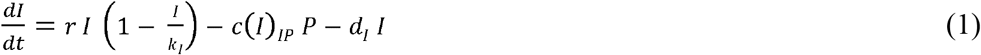

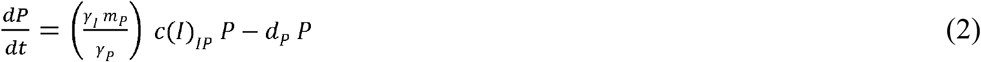

Invertebrates grow logistically, equilibrating in the absence of predators at *k*_*I*_ (1 – *d*_*I*_*/r*). Invertebrates are cleared by predators according to *c(I)*_*IP*_. Clearance *c(I)*_*IP*_ follows a type II functional response (Figure 1A), demonstrated both by sculpins (Mace 1983, Soluk 1993) and round gobies (Laverty et al. 2017, Franta et al. 2023):

**Figure 1.**
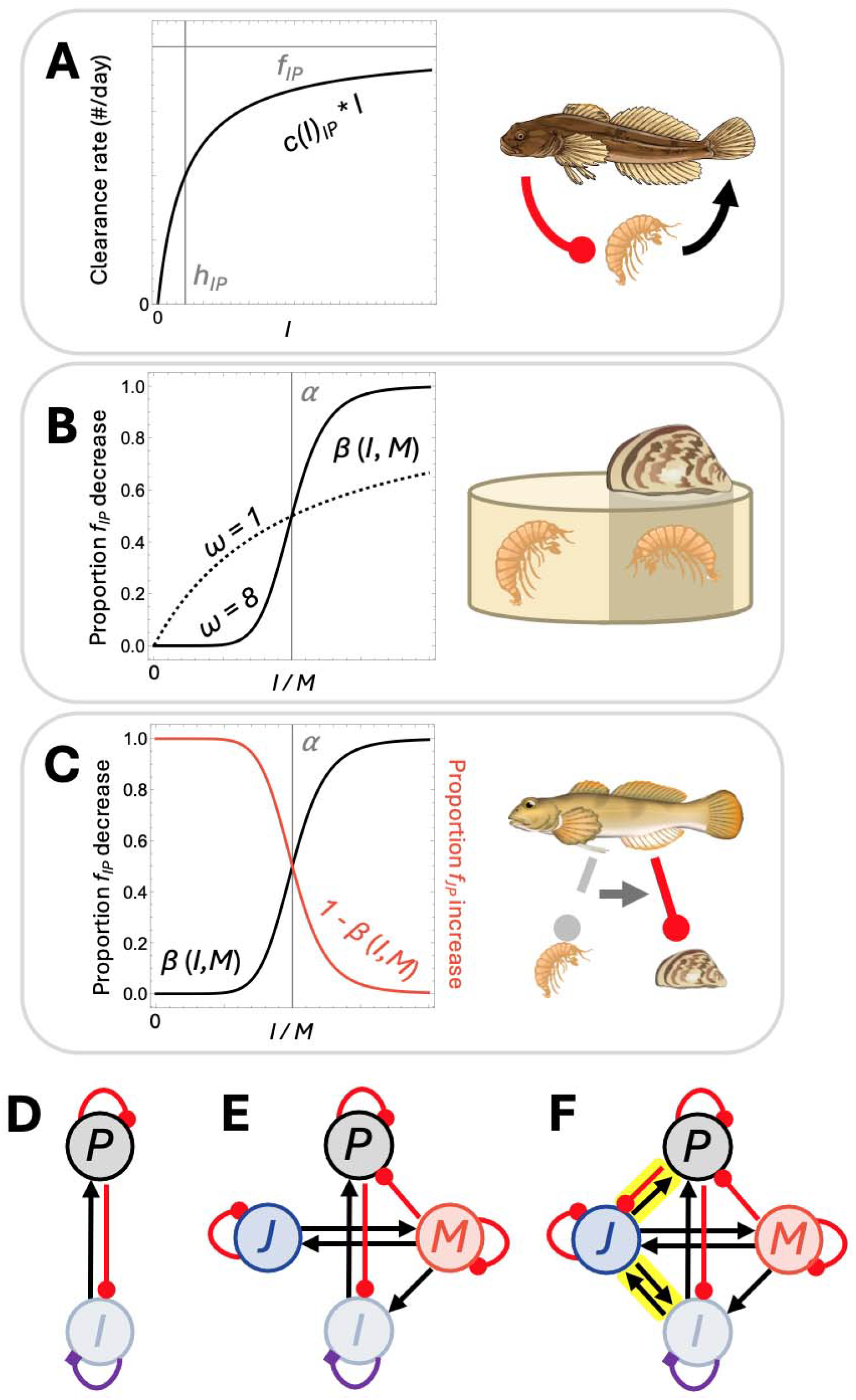
Building blocks of the predator-prey models with and without habitat complexity (HC) and edible HC (EHC). **(A)** The base predator-prey relationship between invertebrates (*I*) and the mid-trophic predator (*P*) follows a type II functional response, where prey cleared per day is a saturating function depending on the maximum feeding rate *f*_*IP*_ and the half-saturation constant *h*_*P*_ where feeding rate is half the maximum. **(B)** HC function, or the influence of *Dreissena* on *P* predation of *I* (scaling *f*_*IP*_). Low invertebrate/mature *Dreissena* (*I/M*) ratios drastically reduce clearance of invertebrates while high ratios render clearance largely the same as without HC. The steepness of this HC shift around threshold level *α* depends on *ω*, with higher values of *ω* yielding a sharper shift. **(C)** With round goby as *P*, foraging shifts from *I* to juvenile *Dreissena* (*J*) symmetrically with HC according to the same *I/M* ratio. **(D - F)** Three benthic food webs examined in this study. Black arrows indicate positive relationship for population growth, such as the positive impact of prey to their predator. Red punches indicate a negative relationship for population growth, such as prey from their predator. The sign of self-limitation of *I* is of particular interest given it varies in sign depending on parameters, indicated by purple diamonds. **(D)** No HC is present in the base *IP* model. *P* benefits from *I* while *I* is inhibited by *P*. The sign of *I*’s self-regulation depends on parameter values. **(E)** With sculpin as *P*, consumption of *I* is mediated by HC from *M. J* and *M* are self-promoting according to births and maturations. **(F)** With round goby as *P*, two new pairwise relationships are created (highlighted in yellow). Round goby (*P*) can now consume *J*, which serve as EHC. Thus, *J* and *I* are mutually promoting, given that an increase in one’s density reduces foraging pressure on the other.

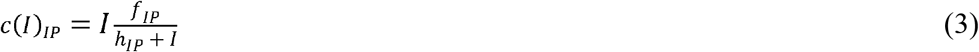

where *f*_*IP*_ is the maximum feeding rate and *h*_*IP*_ describes the invertebrate density where feeding rate is half that maximum. Predators convert cleared invertebrates into new predators according to each taxon’s mass (*γ*_*I*_ and *γ*_*P*_) and predator metabolic efficiency (*m*_*P*_). Invertebrates and predators die at mortality rates *d*_*I*_ and *d*_*P*_, respectively.

### *Dreissena* stage structure

A stage-structure model describes the production of juveniles (*J*) by mature *Dreissena* (*M*) and the maturation of *J*:

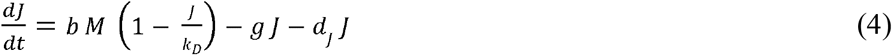

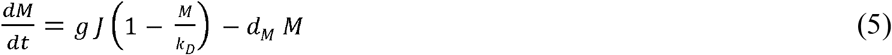

where *b* is the birth rate of *J* by *M, g* is the maturation rate, *k*_*D*_ is the carrying capacity of *Dreissena*, and juvenile and mature *Dreissena* mortality are given by *d*_*J*_ and *d*_*M*_, respectively.

### HC via *Dreissena*

Mature *Dreissena* act as HC providing refuge for *I*. We assumed HC affects maximum feeding rate *f*_*IP*_ (Mocq et al. 2021). Given the threshold hypothesis (Gotceitas and Colgan 1989), we expected changes in *f*_*IP*_ to be non-linear depending on the ratio *I/M*. To explore a variety of potential HC-predation functions (*β*), we used the following form:

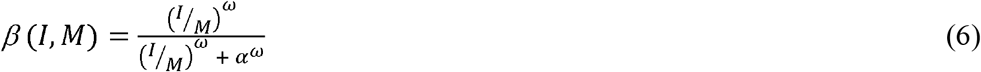

*α* defines the *I/M* ratio at which predation rate is half of its maximum in the absence of HC and *ω* controls the steepness of that change, where *ω* = 1 yields a saturating curve and *ω* > 1 yields a sigmoidal shape (Figure 1B). With HC, clearance of *I* by *P* (equation 3) was scaled by *β*. High *I/M* ratios did not change *f*_*IP*_ (*β(I,M)* = 1) but low *I/M* ratios brought *f*_*IP*_ close to 0 (*β(I,M)* = 0).

Additionally, *Dreissena* increase resource availability for *I* by increasing benthic organic matter (Stewart et al. 1998a, b). With *Dreissena, r* and *k*_*I*_ were scaled by (1 + *δ M/k*_*D*_) where *δ* scales added resources. With sculpin predators, we assumed no other connections between the invertebrate-predator subsystem (equations 1 and 2) and the *Dreissena* subsystem (equations 4 and 5; Figure 1D). However, predation was more complicated with round goby.

### Round goby adaptive foraging

A consumptive link was created between *P* and *J* when round goby was the predator. Round gobies prefer invertebrates but will eat *Dreissena* when invertebrates are not available (Barton et al. 2005, Naddafi and Rudstam 2014). We assumed round gobies switch to consuming *J* symmetrically to the HC benefit for *I* (e.g., van Baalen et al. 2001) according to the inverse of that benefit (1 – *β*; Figure 1C). Thus, predation by round goby was (Fryxell and Lundberg 1994):

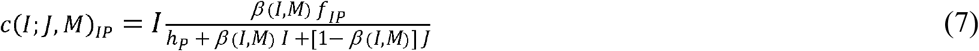

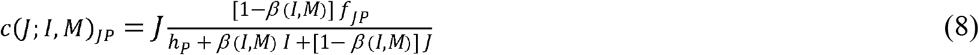

where *f*_*JP*_ is the feeding rate of round goby on *J* and *γ* _*J*_ is the mass of *J*. Total amount of foraging effort in equations 7 and 8 remains equivalent to that in equation 3. Consequently, with the round goby, *Dreissena* acted as a source of EHC, shielding invertebrates from predation and acting as prey when invertebrates were refuged (Figure 1E). The sign structure of the EHC model mirrors those of classic models of omnivory, with juvenile *Dreissena* benefitting both predators and invertebrate prey (Diehl 1992, Vandermeer 2006).

### Simulations

In sum, the final equations describing the three models are:

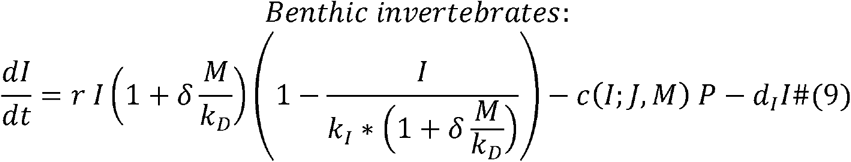

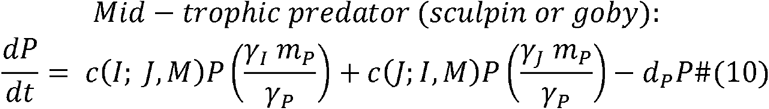

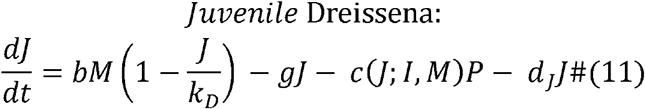

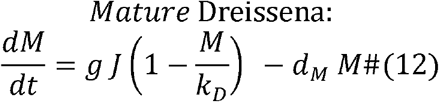

The first model (*IP*) uses only equations 9 and 10 with *J* = 0 and *M* = 0 and the functional response in equation 3. The second model (*IPJM*) uses equations 9 – 12 with *f*_*JP*_ = 0 and the functional response in equation 3. The third model (*IPJM*) uses equations 9 – 12 with *f*_*JP*_ > 0 and the functional response in equations 7 and 8. Values, units, and sources for parameters are in Table S1. First, we analyzed the effect of HC according to predator mortality *d*_*P*_. Then, we compared HC versus EHC according to three parameters: *Dreissena* carrying capacity *k*_*D*_, HC threshold *α*, and HC steepness *ω*. We verified the robustness of our results across three further parameters: *Dreissena* birth (*b*) and mortality (*d*_*J*_) rates and predator conversion efficiency (*m*_*P*_).

In all cases, we examined population densities and the system’s qualitative dynamics (namely, whether population dynamics oscillated). We employed ‘feedback loop tracing’ to determine which biological relationships facilitated and/or eliminated the instability underlying oscillations. Feedback loops are any trophic path in a food web from one species back to itself, where each path is constructed from direct effects in the Jacobian matrix **J** (e.g., *J*_*XY*_ denotes the direct effect of an increase in *Y* on the growth of *X*). Thus, a change in a species’ density can affect its own growth either directly (level 1 feedback, **F**_**1**_) or indirectly via other species (level 2 – 4 feedback, **F**_**2**_ *–* **F**_**4**_). Feedback level *n* aggregates loops of length *n* and products of disjoint loops that contain *n* variables (Figure 2). Stability requires that the sum of each feedback level is negative and that the highest level (**F**_**4**_) is greater than a combination of lower levels (oscillation criterion **OC**; Supplementary section 2; Puccia and Levins 1985). Feedback loop tracing involves dissecting a system to determine each loop’s role in system behavior. Finally, as loop tracing is conducted on equilibria regardless of stability, we demonstrate changes in equilibria in the supplementary material.

**Figure 2.**
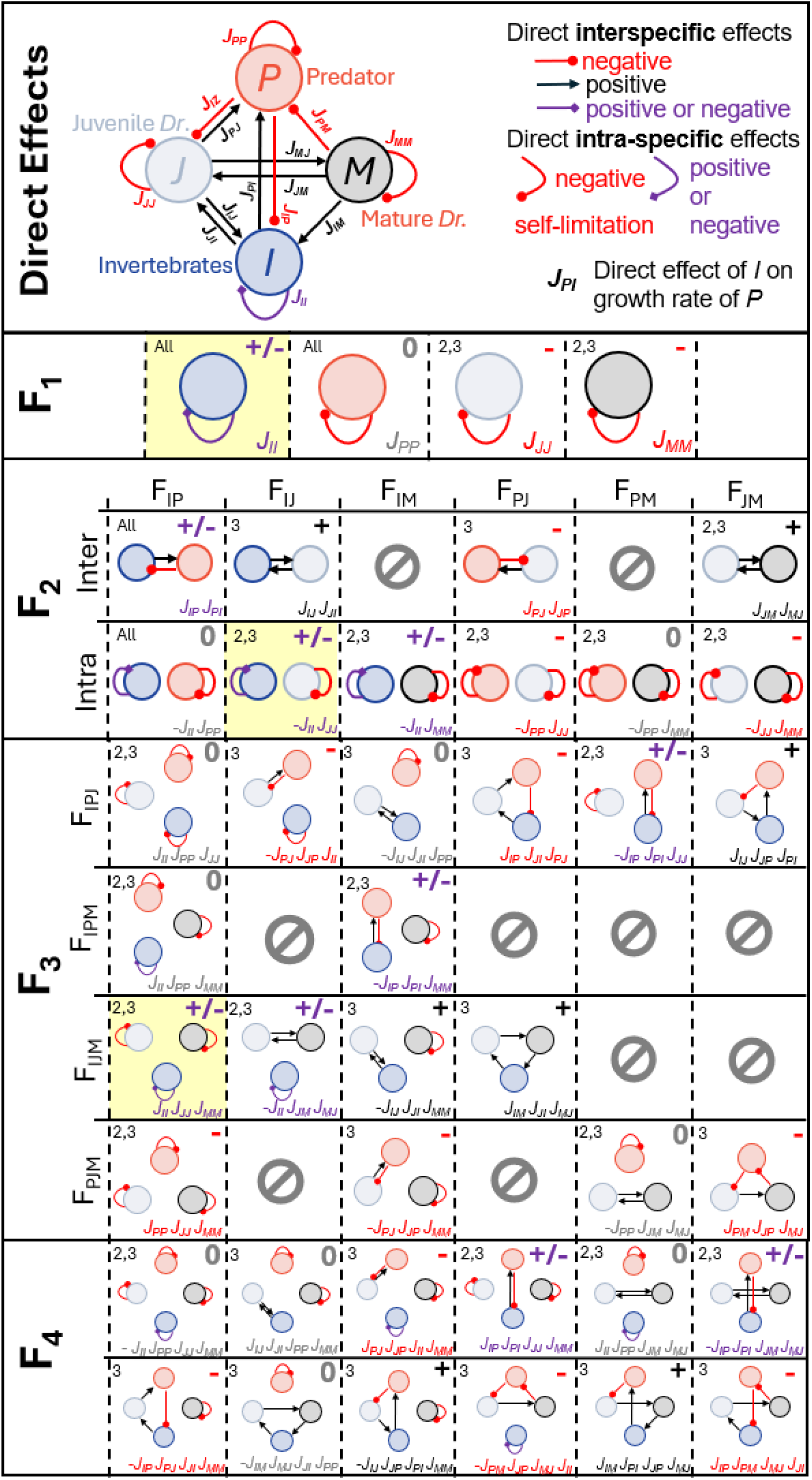
All feedback levels and their constituent loops in these models. Direct effects follow the conventions of Figure 1. Each level of feedback (**F**_**1**_ **– F**_**4**_) is the sum of its constituent feedback loops. Level 2 feedback (**F**_**2**_) is organized according to dyadic intra and interspecific effects. Level 3 feedback (**F**_**3**_) is organized according to each three-species combination. Each feedback loop is provided in its own panel. Relevant models are included in the top left of each panel. The observed sign of each feedback loop is denoted in the top right, with 0 indicating negligible values (<10^-8^). The Jacobian elements that constitute each feedback loop are provided on the bottom right, colored according to the observed sign. Loops of particular emphasis in the results are highlighted in yellow.

## RESULTS

### Model behavior without and with *Dreissena* as HC

The *IP* model oscillated at all predator mortality (*d*_*P*_) except high values where predators persist at low densities (close to *P*_*0*_ = 1 where *P*_*0*_ is the per capita ratio of predator births/deaths; Supplementary section 3). Mean *I* across oscillations increased and equilibrial *I* and *P* both increased with predator mortality before *P* collapsed (Figure 3A, S1A). The introduction of *Dreissena* stabilized a significant range of lower predator mortality, limiting oscillations to high predator mortality and destabilizing the small stable *IP* region. *P* decreased in the stable region, subsequently leveling off in the oscillating region, while *I* increased in both (Figure 3B). Once again, equilibrial *I* and *P* both increased in the oscillating region. Both *I* and *P* were significantly greater with *Dreissena* than without. *Dreissena* densities (*J* and *M*) were constant across predator mortality.

**Figure 3.**
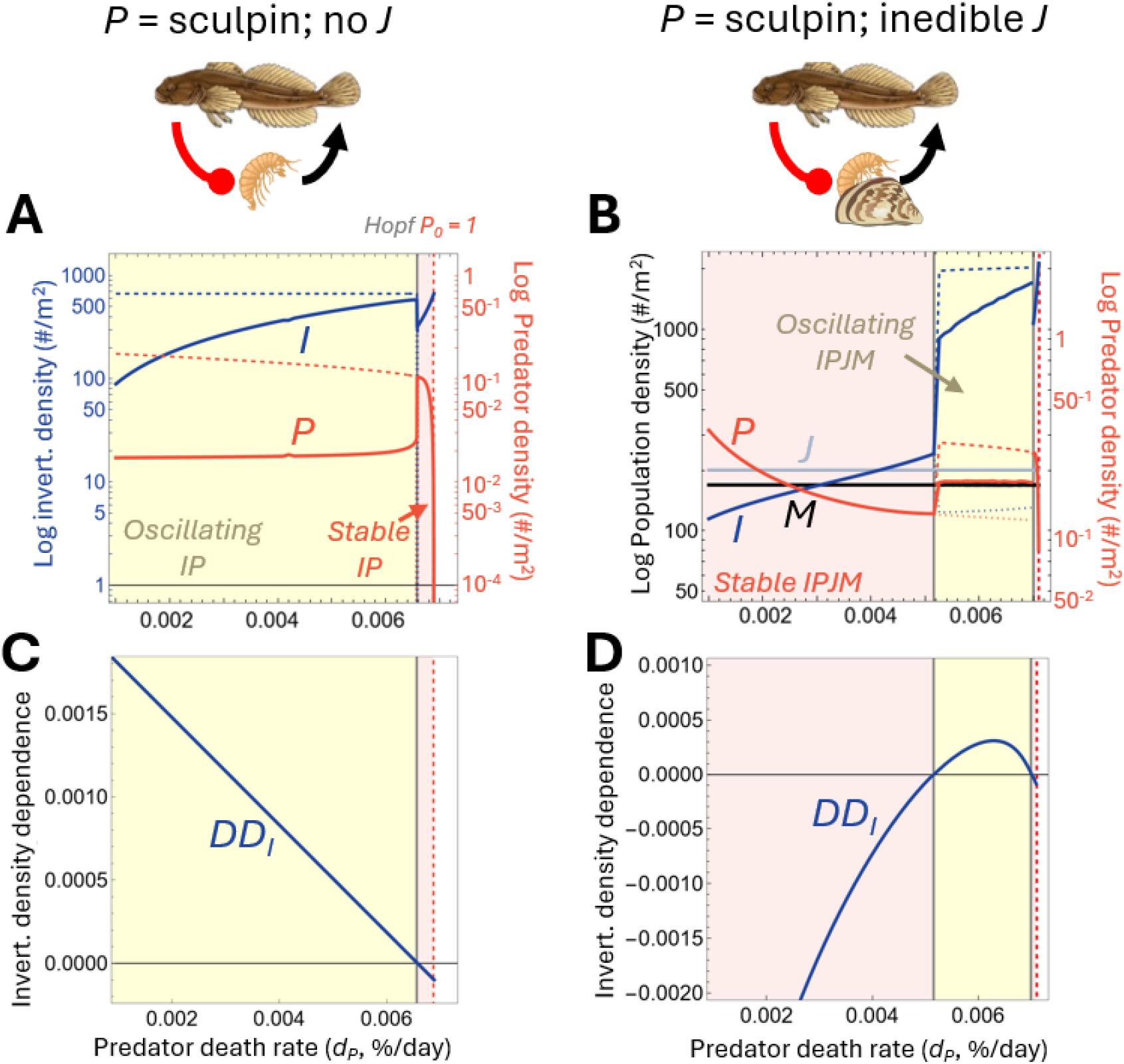
Invading *Dreissena* add habitat complexity (HC) and stabilize predator (*P*) - invertebrate (*I*) dynamics. **(A)** The *IP* model alone has a strong propensity to oscillate (yellow region; solid lines indicate means, dashed lines indicate maxima, and dotted lines indicate minima). Mean invertebrate population increases with predator mortality (*d*_*P*_) until the system briefly stabilizes (Hopf) before the predator population collapses (*P*_*0*_ = 1). **(B)** Juvenile (*J*) and mature (*M*) *Dreissena*, and thus HC, stabilize those predator-prey oscillations at low predator mortality (*d*_*P*_). **(C)** Positive invertebrate density dependence (*DD*_*I*_) at equilibrium 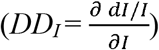 aligns exactly with the oscillating regime without HC. **(D)** *DD*_*I*_ almost exactly aligns with the oscillating region with HC. In all cases, *k*_*D*_ = 200, *α* = 1, and *ω* = 4.

Type II responses destabilize systems by reducing prey density dependence (Gurney and Nisbet 1998, Murdoch et al. 2003). Invertebrate density dependence *DD*_*I*_ is derived from fitness *F*_*I*_:

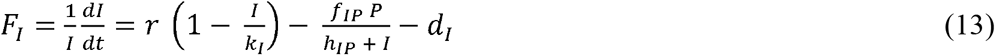

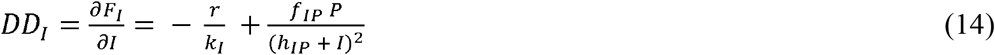

So, *DD*_*I*_ is the sum of two counter-acting forces: self-limitation of population growth due to carrying capacity 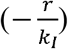 and population clearance from predation 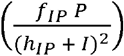. In our models, when self-limitation outweighed clearance, *DD*_*I*_ was negative and dynamics were often stable. When clearance outweighed self-limitation, *DD*_*I*_ was positive and dynamics were often unstable. In the *IP* model, *DD*_*I*_ changed sign exactly at the Hopf that denotes the onset of oscillations (Figure 3C). In the model with *Dreissena, DD*_*I*_ takes a different form:

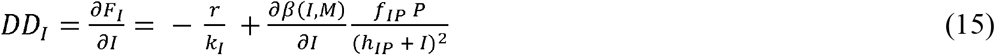

Now, the influence of predation on invertebrate density dependence is scaled by HC via 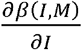, shifting the balance that determines the sign and strength of *DD*_*I*_. As a result, *DD*_*I*_ was negative at lower predator mortality with HC. In this case, invertebrates experienced *‘safety in rarity’* (Murdoch et al. 2003), where an increase in invertebrate density increases their per capita predation risk via skewing towards *I/M* ratios beyond HC threshold *α*.

We used loop tracing to verify the role of *DD*_*I*_ in the onset of oscillations. Notably, feedback loop *J*_*II*_ in **F**_**1**_ (Figure 2) is proportional to *DD*_*I*_:

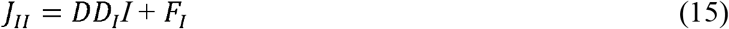

In the *IP* system, **F**_**1**_ was positive in the oscillating region and thus drove those oscillations (Figure 4A). *J*_*II*_ singly accounted for **F**_**1**_’s behavior (Figure 4C). With *Dreissena*, **F**_**1**_ was stabilizing due to high density dependence of *Dreissena* (Figure S2). However, oscillations began according to **OC** driven by **F**_**2**_ and were sustained by positive **F**_**2**_ and **F**_**3**_ (Figure 4B). The combined self-effects of *I* and *J* (*-J*_*II*_ *J*_*JJ*_) shaped **F**_**2**_’s behavior while the addition of the self-effect of *M* (*-J*_*II*_ *J*_*JJ*_ *J*_*MM*_) drove **F**_**3**_ (Figure 4D). Within both these components, *J*_*II*_ underwent sign changes driving the sign change in **OC** (Figure 4E). So, in both systems, the self-regulation of invertebrates drove oscillations in the entire food web. However, with HC, invertebrate self-regulation was realized through its coupling with that of juvenile and mature *Dreissena*.

**Figure 4.**
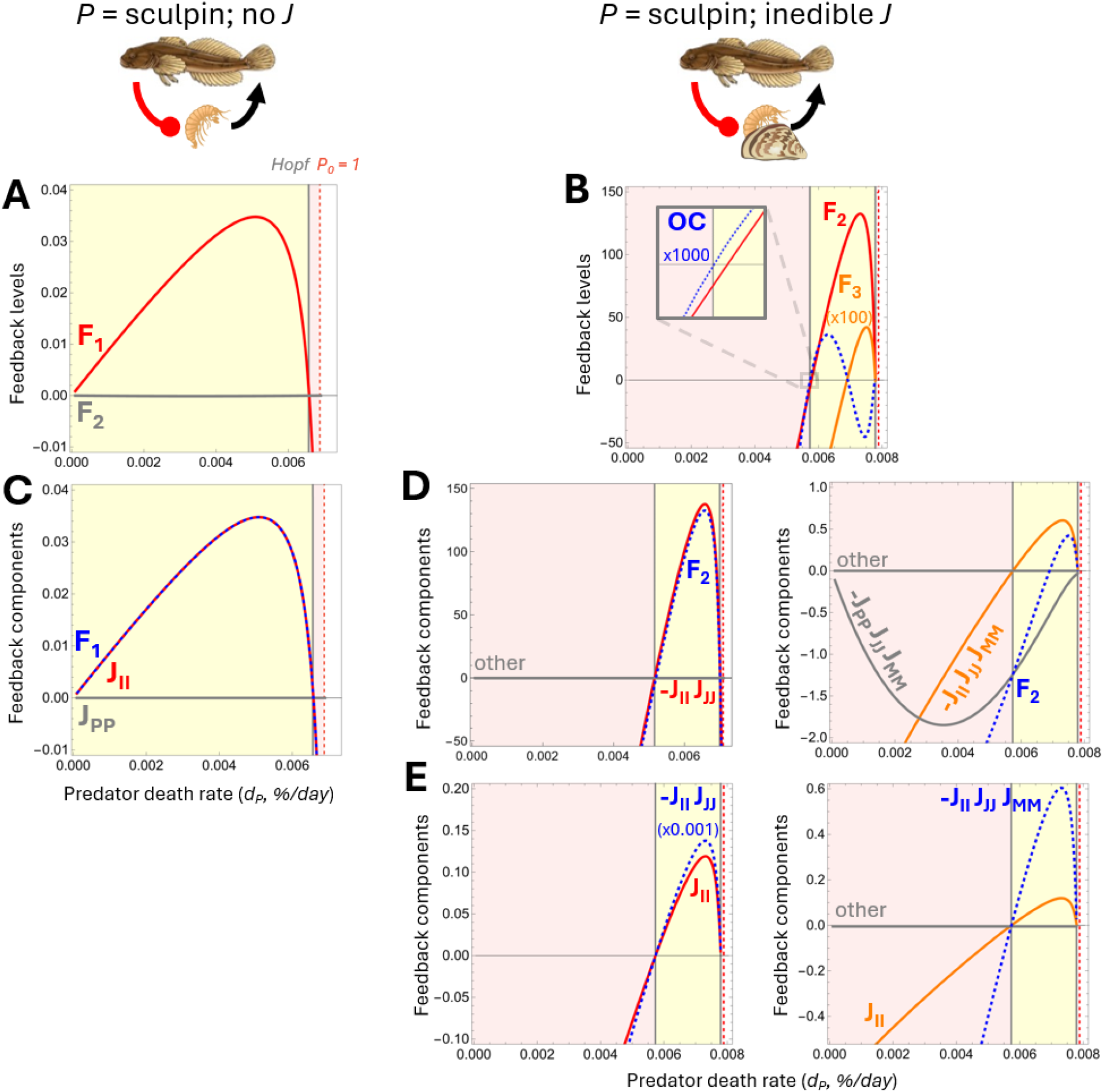
Loop tracing the cause of oscillations in the *IP* and *IPJM* models according to predator mortality *d*_*P*_ with sculpin as the mid-trophic predator *P*. Dashed blue lines indicate the feature to be explained, while red lines indicate the explaining feature (red and gray are aggregated in dashed blue components). **(A)** Without *Dreissena* as habitat complexity (HC), the invertebrate-predator model oscillates when **F**_**1**_ is positive. **(B)** With *Dreissena* (and thus HC), **OC** determines the onset of oscillations (inset) in the invertebrate-predator system. Positive **F**_**2**_, and later **F**_**3**_, drive **OC**’s behavior and continue destabilizing the system after **OC** falls below 0. **(C)** Without HC, the self-effect of invertebrates **J**_**II**_ drives the behavior of **F**_**1**_. **(D)** With HC, the self-effect of invertebrates scaled by the self-effect of juvenile *Dreissena* -*J*_*II*_ *J*_*JJ*_ explains almost all the behavior of **F**_**2**_ (left) while the added self-effect of mature *Dreissena* (yielding **-***J*_*II*_ *J*_*JJ*_*J*_*MM*_) explains almost all the behavior of **F**_**3**_ (right). Destabilization of **F**_**3**_ lags behind **F**_**2**_ because of the inverse effect of *Dreissena* amplifying the self-effect of predators (*-J*_*PP*_ *J*_*JJ*_*J*_*MM*_). **(E)** Finally, in both **F**_**2**_ (left) and **F**_**3**_ (right), self-regulation of invertebrates *J*_*II*_ corresponds with relevant sign changes and thus is primarily responsible for oscillations in the *IPJM* system. In all cases, *k*_*D*_ = 200, *α* = 1, and *ω* = 4.

### Model behavior with HC v. EHC

To examine model behavior with HC versus EHC across HC-predation functions, we held predator mortality constant (*d*_*P*_ = 0.004; unstable *IP*, stable *IPJM*) and varied *Dreissena* carrying capacity (*k*_*D*_), HC threshold (*α*), and HC steepness (*ω*). With HC, each of *I, J*, and *M* increased with *Dreissena* carrying capacity while *P* first increased then decreased, leading to extirpation (Figure 5A). With EHC, *I, J*, and *M* still increased but were significantly reduced while *P* increased at much higher densities (Figure 5B).

**Figure 5.**
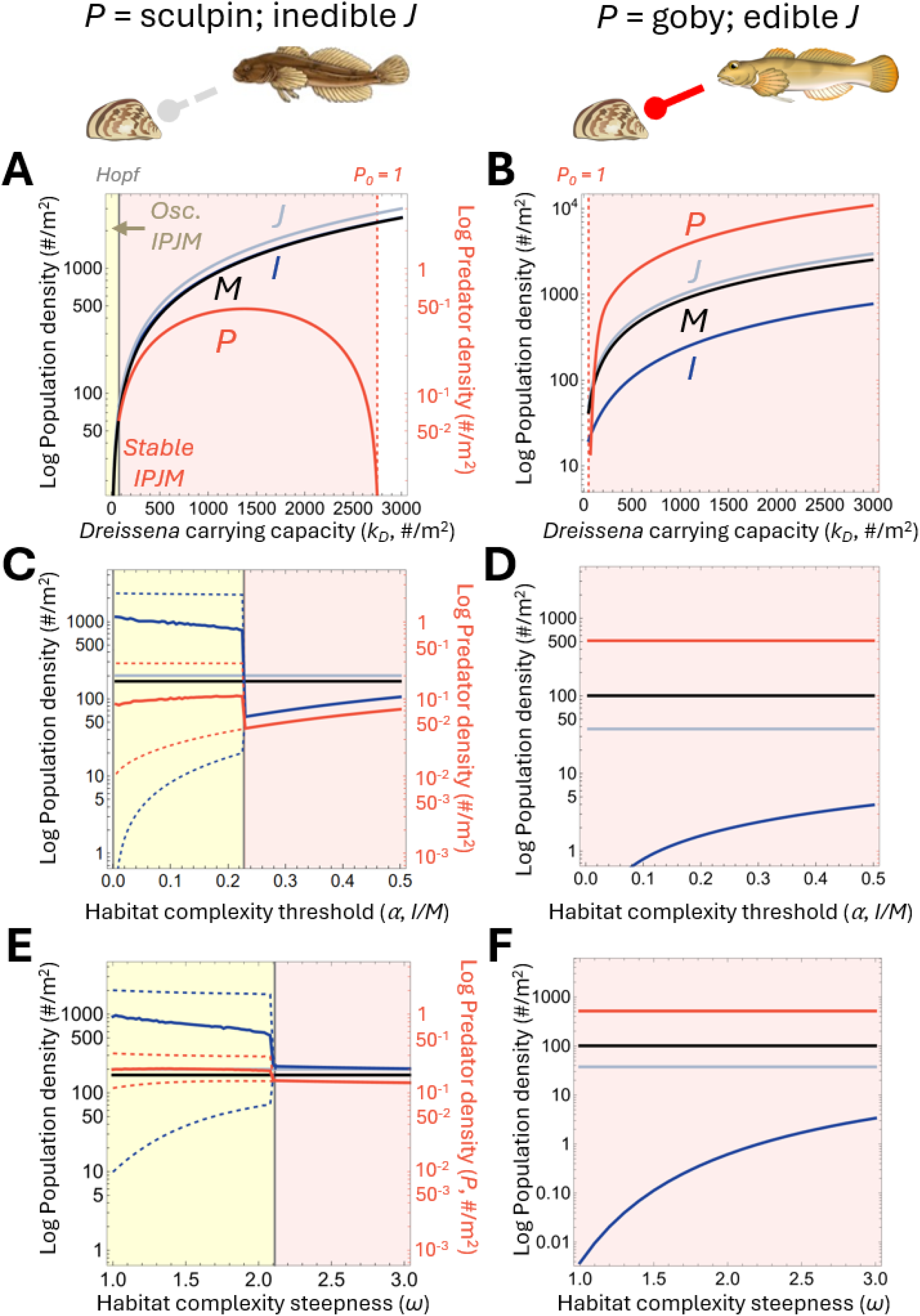
EHC further stabilizes predator-prey dynamics. Conventions follow those in Figure 3. **(A)** Increasing the carrying capacity of *Dreissena k*_*D*_ increases habitat complexity (HC) and subsequently leads to the extirpation of the mid-trophic predator *(P*; *P*_*0*_ = 1). **(B)** When juvenile *Dreissena* (*J*) are edible (thus, edible habitat complexity ‘EHC’), those juveniles become fixed across their carrying capacity and *Dreissena* productivity is translated into increasing predator density. Both invertebrates (*I*) and mature *Dreissena* (*M*) continue to increase given that mature *Dreissena*, which confer HC, are not themselves edible and can still accrue larger populations with increasing *k*_*D*_ from the fixed juvenile population. **(C)** Even with HC, the predator-invertebrate model can still oscillate (yellow region) when the HC threshold *α* drops. **(D)** With EHC, oscillations disappear. Invertebrates and both juvenile and mature *Dreissena* are held at a significantly lower densities while the predator density is significantly greater. **(E)** Reducing HC threshold steepness *ω* can also yield oscillations without EHC. **(F)** Similarly to changes with HC threshold *α*, oscillations according to HC steepness *ω* disappear with EHC. In all cases, *d*_*P*_ = 0.004, *k*_*D*_ = 200, *α* = 1, and *ω* = 4 unless varied.

Although both models with HC and EHC were generally stable across *Dreissena* carrying capacity, HC model stability changed with HC threshold *α* and steepness *ω*. With HC, decreasing HC thresholds first decreased *I* and *P*, then destabilized the system, creating increasingly large oscillations with greater mean *I* and *P* (Figure 5C). Equilibrial *I* and *P* both decreased in the oscillating region (Figure S3). Using loop tracing as above (Figures S4, S5), we found that **F**_**2**_ drove the behavior of **OC**, destabilizing the system at low HC thresholds. Once again, the self-regulation of invertebrates drove **F**_**2**_’s behavior by its coupling to the self-regulation of juvenile *Dreissena* (*-J*_*II*_ *J*_*JJ*_). Conversely, with EHC, *J* and *M* were fixed at lower densities, *P* was fixed at a much higher density, and *I* was fixed at the lowest density, slowly increasing with the HC threshold. The weakening of **F**_**2**_ with a decreasing HC threshold was eliminated with EHC, as both invertebrate and juvenile dreissenid self-regulation were held relatively fixed. So, HC could stabilize the *IP* system at some HC thresholds, while EHC stabilized the system across all thresholds.

Densities and stability also depend on HC steepness (*ω*). In the HC model, steeper predation rate changes with changing *I/M* yielded stable dynamics while gradual predation rate changes yielded oscillations (Figure 5E, F). More gradual HC functions increased *I* and the amplitude of oscillations, while both equilibrial *I* and *P* were greater with more gradual HC functions (Figure S6). The change to EHC yielded similar results for HC steepness as the HC threshold. *P* was higher, *I* was lower, and *J* and *M* were fixed at lower values. Oscillations with gradual HC functions, still driven by *-J*_*II*_ *J*_*JJ*_ within **F**_**2**_, were eliminated with EHC (Figure S7). Once again, HC could stabilize the *IP* system at some HC steepness, while EHC stabilized the system across all steepness.

Finally, we tested if our results held across *Dreissena* birth (*b*) and mortality (*d*_*J*_) rates and predator conversion efficiency (*m*_*P*_). Neither *Dreissena* birth or mortality rate impacted the qualitative dynamics or the equilibria of each model, instead only affecting the rate upon which those equilibria converge. Predator conversion efficiency did alter stability in the HC model via changing predator populations and thus the clearance component of *DD*_*I*_. However, the introduction of EHC once again quashed any oscillations across the tested conversion efficiency range (Figure S8).

## DISCUSSION

We used our models to explore when and why HC and EHC change population dynamics and benthic food web stability. Our models predicted higher predator densities with EHC, consistent with observed differences between sculpin and round goby in *Dreissena*-infested waters (Nalepa et al. 2009). Round goby benefited from the additional available prey item (juvenile *Dreissena*) but also from the loss of refuge for their preferred invertebrate prey (Diggins et al. 2002; Foley et al. 2017). In natural systems, *Dreissena* are not extirpated where round gobies invade (Madenjian et al. 2010) despite round goby’s ability to clear large amounts of *Dreissena* (Naddafi and Rudstam 2014). Instead, *Dreissena* size distributions skew larger at depths with round gobies (Karatayev et al. 2023). Our models captured both the persistence and changed size distribution of *Dreissena* (*J* v. *M*) with round goby. Additionally, our models suggested that edibility of *Dreissena* was more detrimental for invertebrate populations than *Dreissena* populations. Increases in invertebrate densities with HC via *Dreissena* and subsequent decreases in invertebrate densities with EHC via round goby in these models have been observed empirically (Kuhns and Berg 1999, Kipp and Ricciardi 2012, Brooking et al. 2022). As such, in a broader meta-community we expect invertebrates to be most abundant with abiotic HC, followed by EHC, followed by exposed areas.

Additionally, our models recapitulated results suggesting abiotic HC can stabilize predator-prey interactions (Huffaker 1958, Scheffer and De Boer 1995, van Nes and Scheffer 2005). However, using loop tracing, we were able to show *why* HC is stabilizing. The self-regulation of prey (*J*_*II*_) underlies stability changes (Oaten and Murdoch 1975, Murdoch et al. 2003, Forbes and Hall 2025). HC stabilizes predator-prey interactions by providing safety in rarity for prey (thus negative *J*_*II*_). Only at predator mortality where invertebrate densities significantly exceeded those of *Dreissena* did our model with HC still oscillate. Additionally, sometimes HC can shift type II responses to type III responses (e.g., Barrios-O’Neill et al. 2016). Self-regulation of prey determines stability for type III responses as well (Forbes and Hall 2025). So, whether HC changes the form of a functional response (e.g., Lipicius and Hines 1986; Alexander et al. 2012) or not (e.g., Mocq et al. 2021), self-regulation of prey appears to drive stability.

EHC also had profound impacts on the behavior and stability of our modeled benthic food webs. With just HC, low HC thresholds and steepness (more gradual changes to predation) were destabilizing. Across examined HC thresholds and steepness, the switch from HC to EHC stabilized the food web via increased predator densities strengthening both invertebrate (*J*_*II*_) *and Dreissena* (*J*_*JJ*_, *J*_*MM*_) self-regulation. Stabilizing EHC aligns with existing work suggesting prey-switching is stabilizing (Fryxell and Lundberg 1994, Křivan 1996, Post et al. 2000, van Baalen et al. 2001). In our case, we were able to demonstrate *why*: prey switching with EHC provided safety in rarity for both prey species. That said, not all prey-switching functions are stabilizing (e.g., Oaten and Murdoch 1975, Matsuda et al. 1986) and other switching functions with EHC deserve exploration. Additionally, we recommend further exploration of EHC in relation to prey guilds (Diehl 1992), where certain members of a guild may benefit from EHC and guild members may compete for access to EHC.

We focused on HC’s influence on predator feeding rates, but other HC mechanisms (e.g., limited prey mobility; Boukal et al. 2007) warrant further exploration. Nevertheless, our models aid progress to better understand HC and contribute to understanding trophic interactions that depend on several species (Abrams 2022). Finally, loop analysis (Puccia and Levins 1985) and loop tracing serve as tools to move from description of model behavior to their mechanisms. Loop tracing serves not only the analysis of theoretical models but is also a powerful comparative tool for natural communities (e.g. Scotti et al. 2020, Capelli et al. 2025). Dissecting biological feedback in food webs will be especially important for understanding trophic interactions that depend on several species, including those that play unique roles such as EHC.

## ACKNOWLEDGEMENTS

This project was supported by Award 2023-MAR-95003 from the Great Lakes Fishery Commission. Thank you to the Rubenstein Ecosystem Science Laboratory and two anonymous reviewers for their feedback on this manuscript.

## SUPPLEMENTARY MATERIAL

### Section 1: Model parameters

**Table S1:**
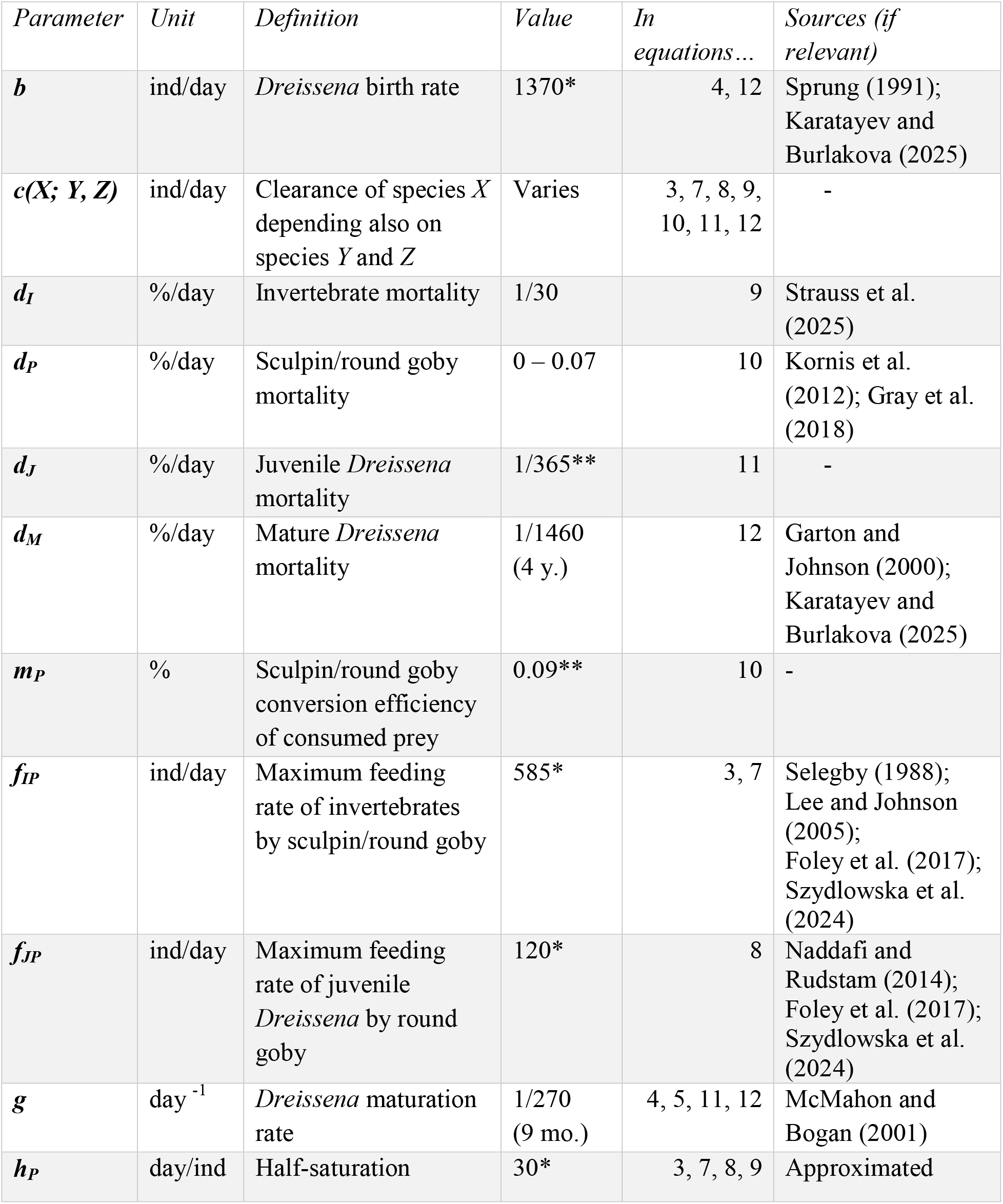
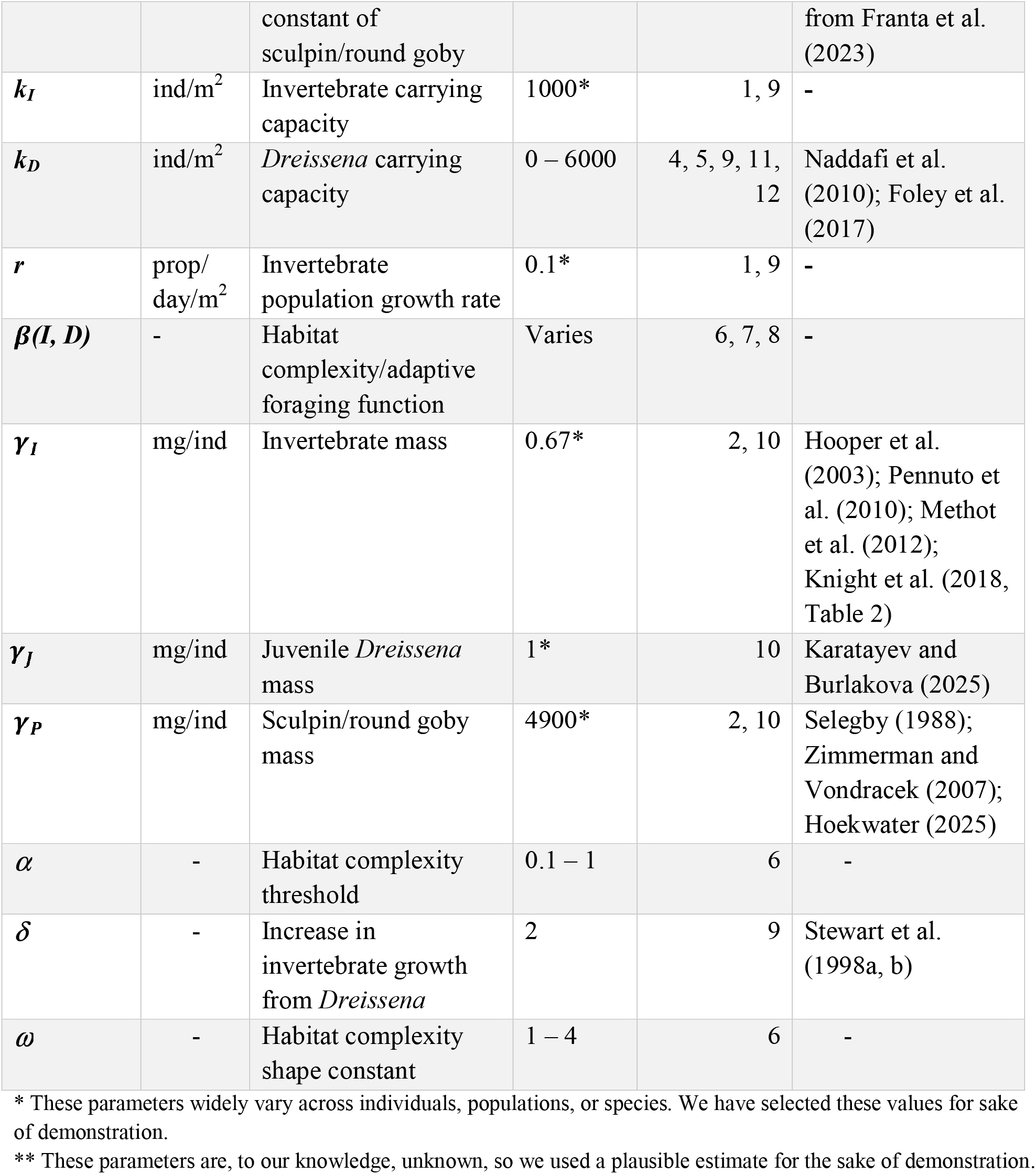
Model parameters, units, and sources.

### Section 2: Feedback loops from the Jacobian matrix

Feedback loops are described by a system’s characteristic polynomial and Jacobian matrix, where Jacobian element *J*_*XY*_ notes the direct effect of an increase in density *Y* on growth rate of *X* (i.e., *J*_*XY*_ = *∂G*_*X*_/*∂Y*; Puccia and Levins 1985). A feedback loop is any assemblage of these direct effects that lead a species back to itself. Feedback loops thus can be any link from 1 to *N*, where *N* is the dimensionality of the food web.

Our arguments regarding the role of HC, EHC, and stability hinge on the use of feedback loops. The decomposition of those various loops during ‘loop tracing’ depends on first specifying Jacobian matrices (**J**) of each model. These matrices describe the direct effects of each species on themselves or on each other. The feedback loops, connecting species here in loops of lengths one (**F**_**1**_), two (**F**_**2**_), three (**F**_**3**_), or four species (**F**_**4**_), organize those direct effects into paths of indirect effects. We have demonstrated these loops pictorially in Figure 2 of the main text. Here, we write out in full the value of each loop and its situation in each Jacobian matrix.

#### (A) Jacobian matrices for each model

Each model has its own Jacobian matrix depending on the biological assumptions codified into those models. First, we consider the Jacobian of the predator-prey model without *Dreissena* (**J**_**V1**_, main text equations 1 and 2). That Jacobian is (in order of *I* and *P*):

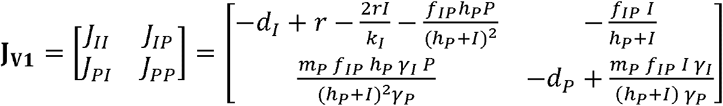

where element *J*_*XY*_ is the effect of an increase in density of *Y* on the growth rate of *X, G*_*X*_ (so **J**_**XY**_ = *∂G*_*X*_/*∂Y*). The addition of *Dreissena* adds dimensions and new terms to the Jacobian (main text equations 10 – 13). The Jacobian with *Dreissena* (**J**_**V2**_) is (in order of *I, P, J*, and *M*):

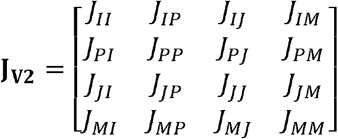

In the case of inedible *Dreissena* (sculpin as mid-trophic predator, *f*_*JP*_ = 0):

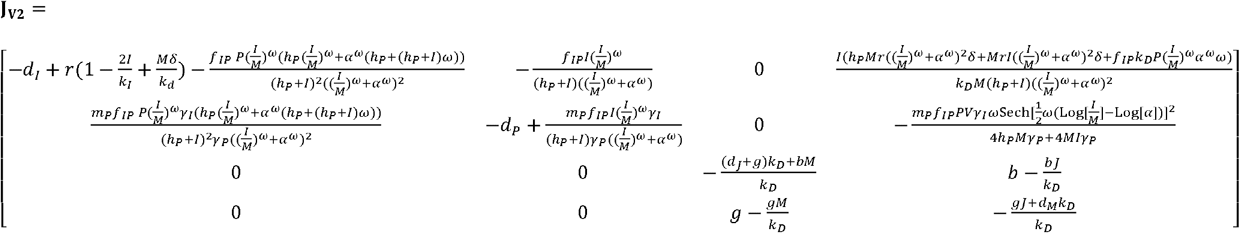

The matrix re-affirms the lack of direct relationships between the predator-prey model and the *Dreissena* stage-structured model, with only the influence of *M* on *I* (*J*_*IM*_) and *P* (*J*_*PM*_) taking on non-zero values (while all other potential direct relationships hold 0 value).

With edible *Dreissena* (goby as *P, f*_*JP*_ > 0) **J**_**V2**_ changes substantially. Here, for the sake of space, we present **J**_**V3**_ sequentially:

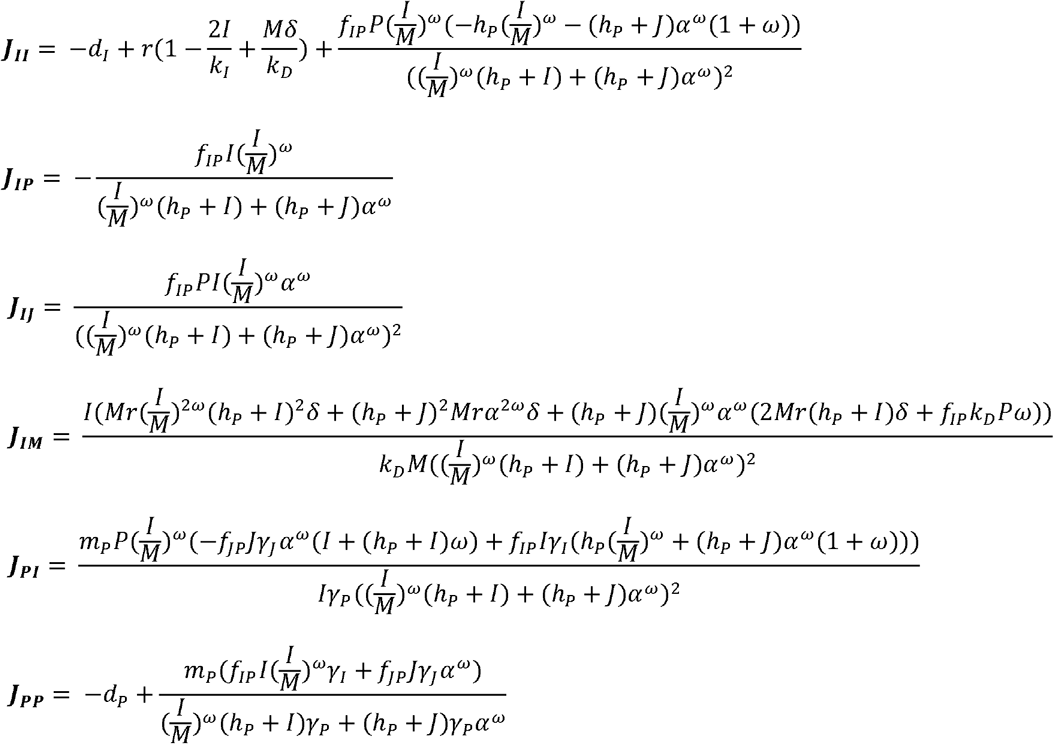

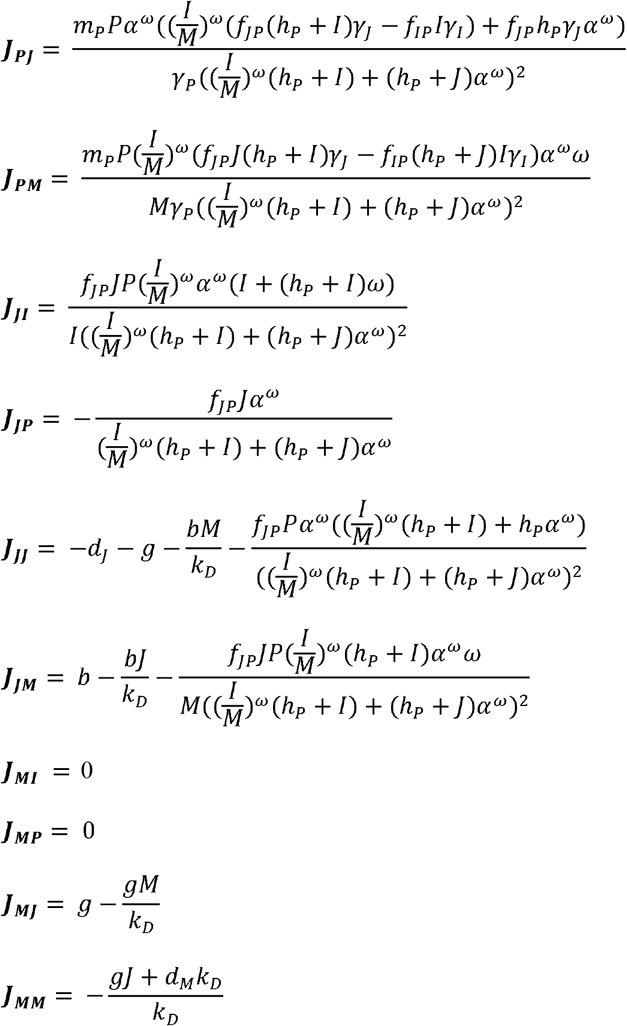

In this case, only the direct effect of *I* on *M* (**J**_**MI**_) and the direct effect of predators on *M* (**J**_**MP**_) take on 0 value.

#### (B) Feedback loops and criteria for oscillations

Two general conditions determine the onset of oscillations in food webs. First, each level of feedback must be negative. Second, the highest level of feedback must be greater than some combination of lower levels of feedback.

With only two species, oscillations begin when level 1 feedback (*F*_1_) becomes positive (Puccia and Levins 1985). Thus, the *IP* system is stable when *F*_*1*_ < 0. In the main text, we demonstrate how *J*_*II*_ makes **F**_**1**_ positive in oscillating regions. The *IPJM* system has more levels of feedback, thus these criteria change. The *IPJM* system is stable when (Puccia and Levins 1985):

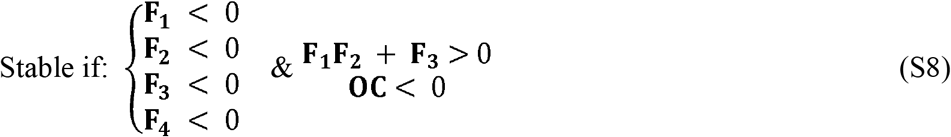

where **OC** is the oscillation criterion. For a food web with four species, **OC** is:

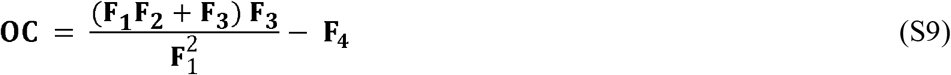

The behavior of **OC** can thus be compared with the behavior of its constituent feedback levels, as demonstrated in the main text.

### Section 3: Invasion of the mid-trophic level predator in these models

‘Transcritical’ bifurcations denote the invasibility or ability to sustain a population for mid-trophic predators in these models. Transcritical bifurcations can be detected by determining whether a population’s growth rate is positive or negative at the equilibrial populations of the existing species. Here, we examine whether the mid-trophic predator can persist at a given parameter value by determining whether its population gains from consumption are greater than its losses from mortality (here using the consumer-resource model [equations 1 and 2] to demonstrate):

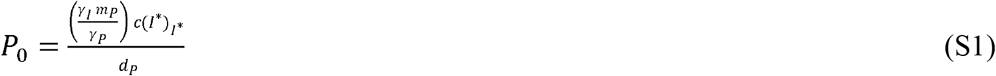

P0 is the reproductive ratio, or the per capita ratio of births to deaths, of predators. When *P*_*0*_ > 1, predator births exceed their mortality, and the population grows. *P*_*0*_ < 1 means predator population will decline until extirpation. When gains from consumption (numerator) are equal to losses from mortality (denominator), the mid-trophic predator can invade the system and maintain a positive density (at *P*_*0*_ = 1). Equation S1 is evaluated at the current densities of existing populations in the model denoted by * (in this case, *I* = k*_*I*_ *(1 – d*_*I*_*/r)* in the absence of predation). Similar criteria can be evaluated for other species in the model, but here only *P* was relevant.

### Section 4: Additional figures demonstrating model and feedback loop behavior for each focal parameter

Here, we provide supplemental figures for population density, equilibrial density, and feedback loop changes with the focal parameters of this manuscript. The conclusions from these analyses are summarized in the main text, while details can be found in the supplemental figures and their captions. While the main text provided plots for mean, maximum, and minimum population densities and the critical feedback loops, the figures below demonstrate both (1) equilibrial densities that are used in feedback loop analysis and (2) all possible feedback loops at each level (see also Figure 2 of the main text). Figures are ordered according to their discussion in the main text.

**Figure S1:**
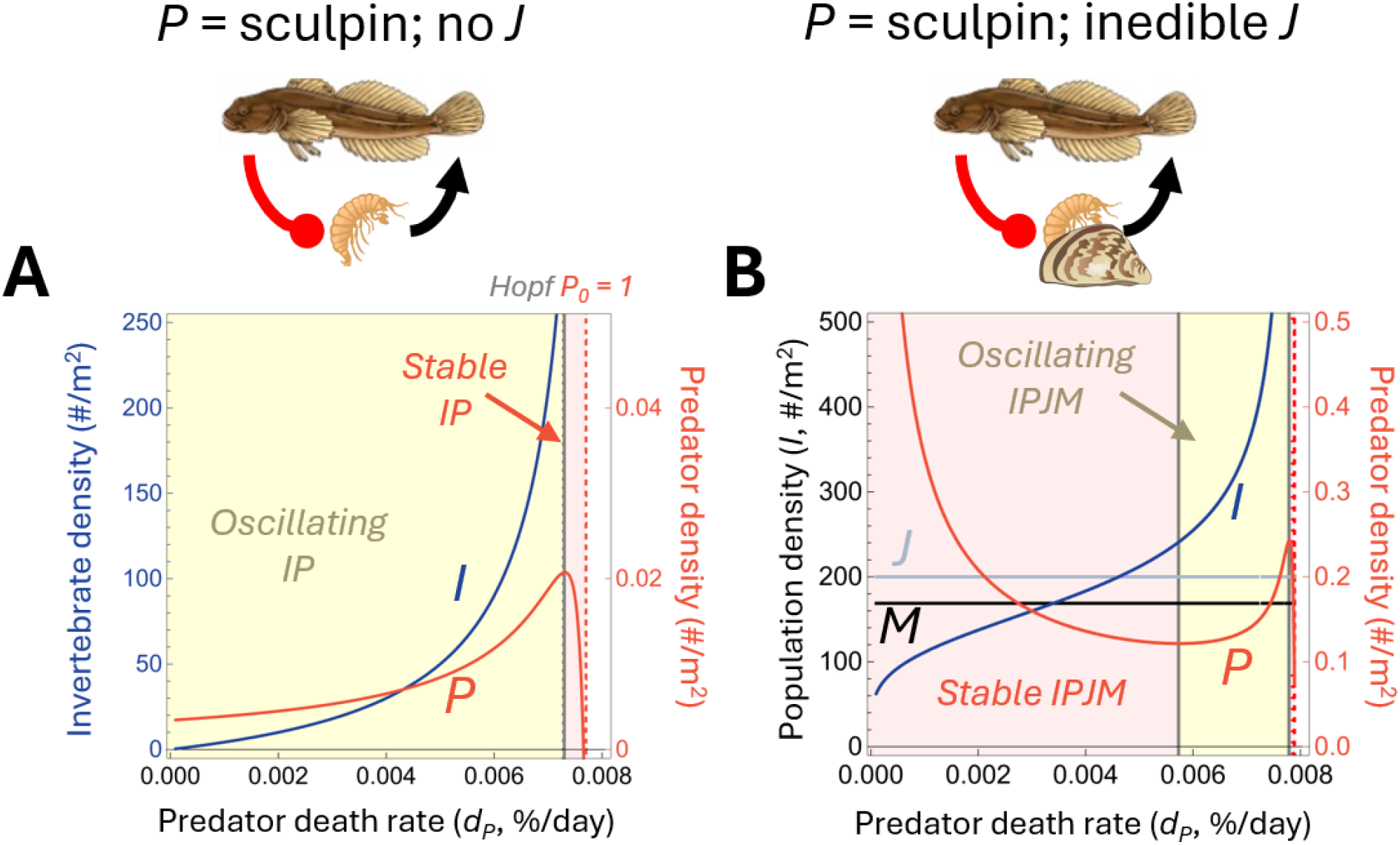
Equilibrial densities for the predator-prey **(A)** and HC **(B)** models according to predator mortality *d*_*P*_. Yellow areas indicate oscillating dynamics while orange areas indicate stable dynamics. The vertical gray line denotes the Hopf bifurcation at the onset of oscillations, while the dashed orange line indicates the invasion criterion for the predator (*P*_*0*_ = 1). This figure corresponds directly to Figure 3 in the main text. In all cases, *k*_*D*_ = 200, *α* = *1*, and *ω* = 4.

**Figure S2:**
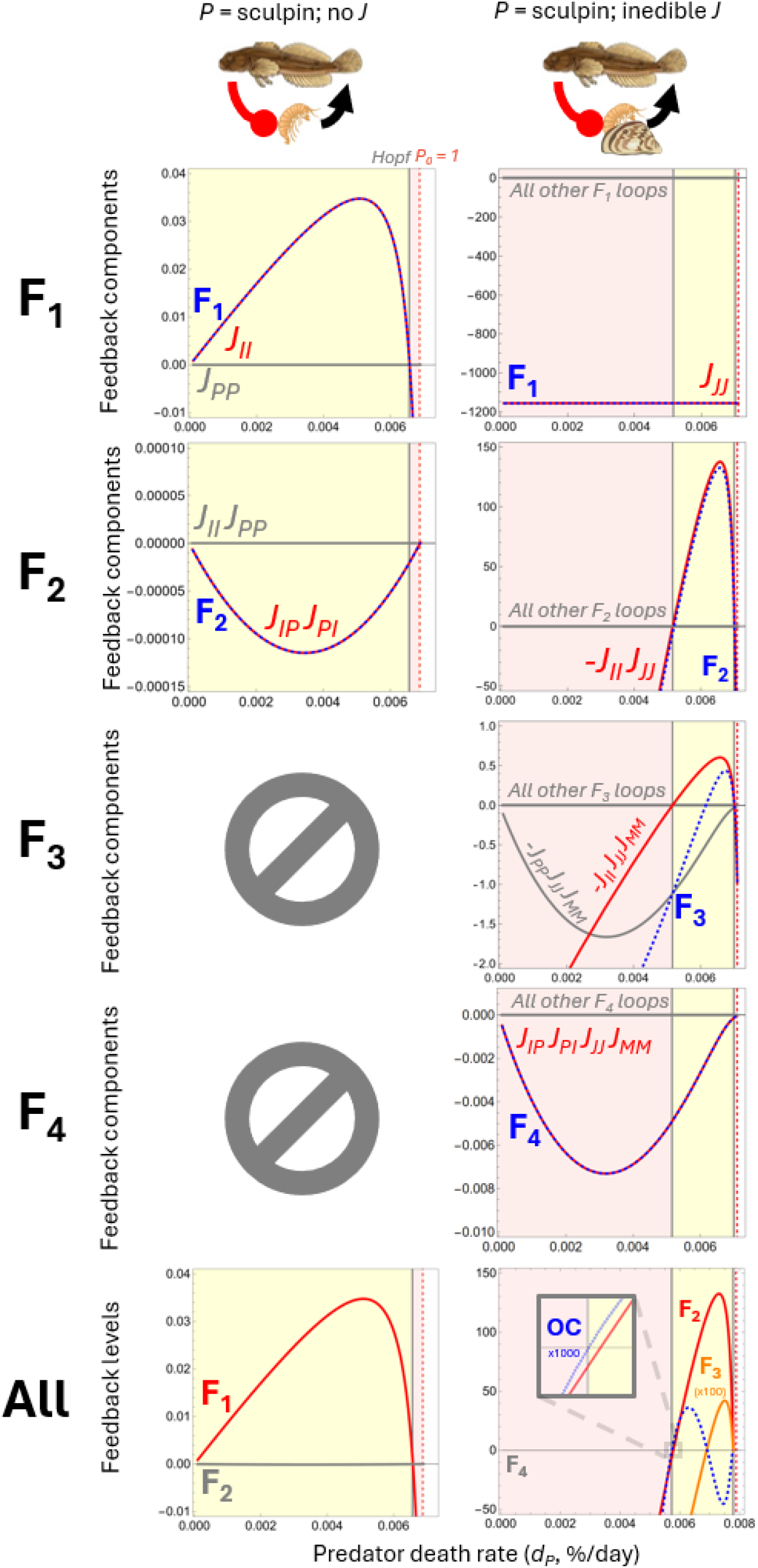
All feedback loops in the predator-prey and HC models according to predator mortality *d*_*P*_. As in the main text, dashed blue lines indicate the feature to be explained, while red/orange lines indicate the explaining feature (so, red and gray components combine to create the dashed blue component). Each row indicates a different level of feedback, with the dashed blue line showing the behavior of the entire level and the other lines describing the behavior of components of that level (see Figure 2 of the main text). This figure corresponds directly to Figure 4 of the main text. In all cases, *k*_*D*_ = 200, *α* = 1, and *ω* = 4 unless varied.

**Figure S3:**
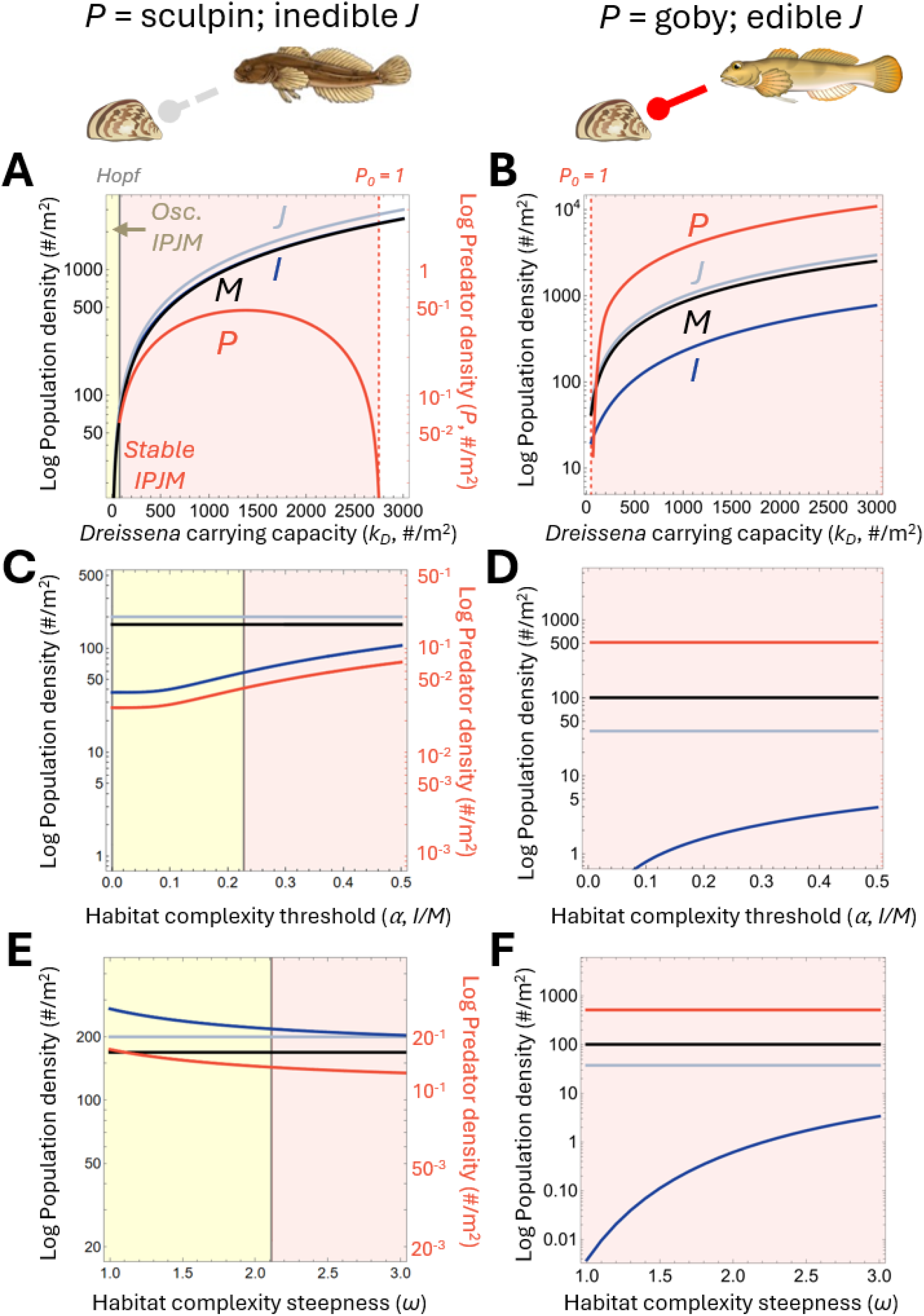
Equilibrial densities for the HC and EHC models according to *Dreissena* carrying capacity **(A & B)**, HC thresholds *α* **(C & D)**, and HC steepness *ω* **(E & F)**. Color conventions are the same as in Figure S1. This figure corresponds directly to Figure 5 of the main text. In all cases, *d*_*P*_ = 0.004, *k*_*D*_ = 200, *α* = 1, and *ω* = 4 unless varied.

**Figure S4:**
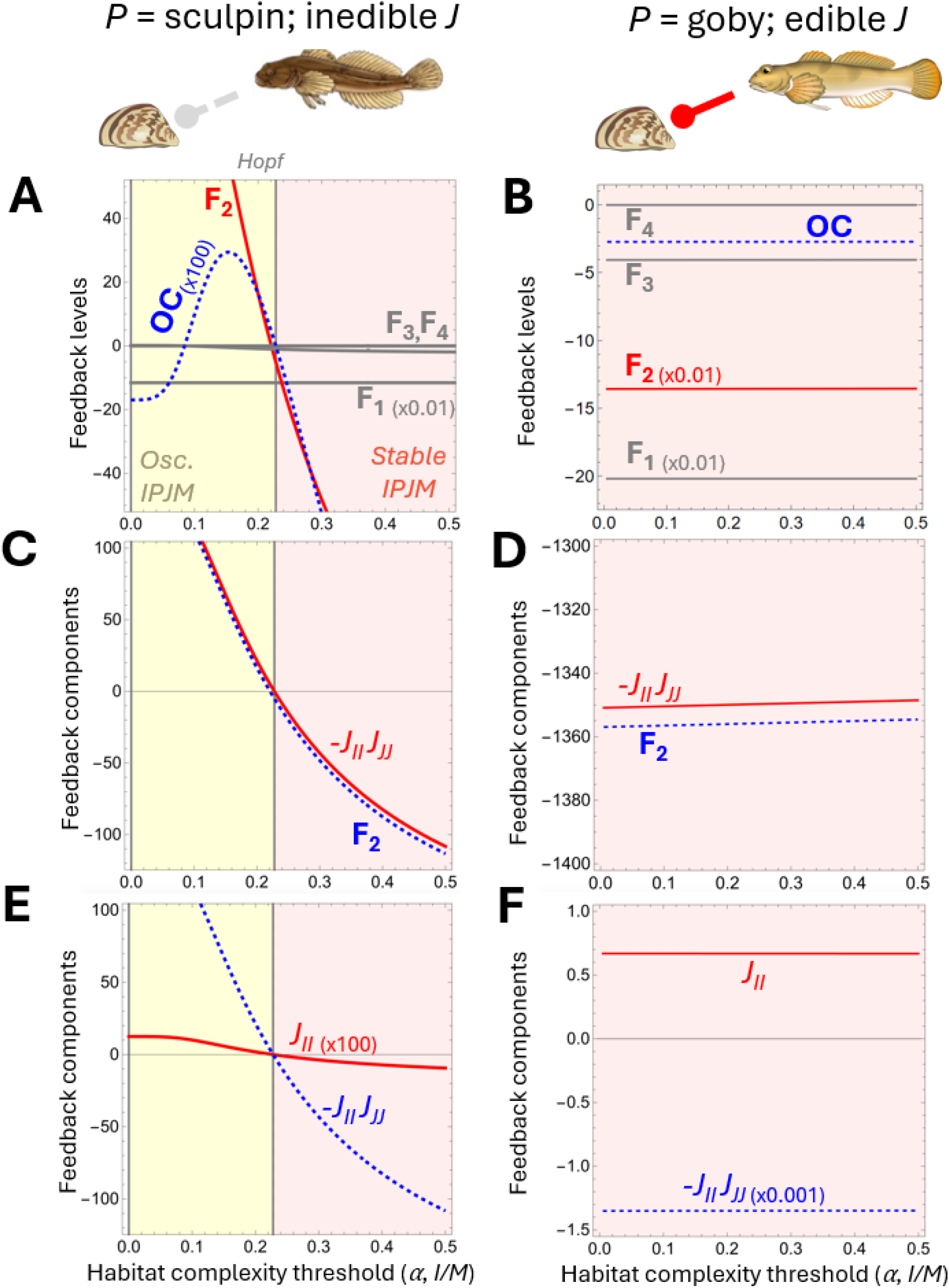
Loop tracing the onset and elimination of oscillations in *IPJM* models with sculpin and round goby as the mid-trophic level predator according to habitat complexity (HC) threshold *α*. Conventions follow Figure 4. **(A)** Without edible habitat complexity (EHC), the food web oscillates according to the oscillation criterion (**OC**) close to when **F**_**2**_ is positive. **(B)** With EHC, feedbacks remain largely unchanged with changing *α*. Neither **OC** or **F**_**2**_ shifts or changes sign as without EHC. **(C)** Without EHC, the self-effect of invertebrates *J*_*II*_ scaled by the self-effect of juvenile *Dreissena J*_*JJ*_ drives the behavior of **F**_**2**_, thus driving oscillations in the invertebrate-predator system. **(D)** With EHC, while *J*_*II*_ scaled by *J*_*JJ*_ does change slightly with *α*, it is extremely stabilizing at all examined values. **(E)** Without EHC, *J*_*II*_ specifically undergoes the sign change relevant to food web stability, thus positive *J*_*II*_ drives oscillations without EHC. **(F)** With EHC, *J*_*II*_ is positive and therefore destabilizing but does not change and is outweighed by other loops in the food web. In all cases, *d*_*P*_ = 0.004, *k*_*D*_ = 200, *α* = 1, and *ω* = 4 unless varied.

**Figure S5:**
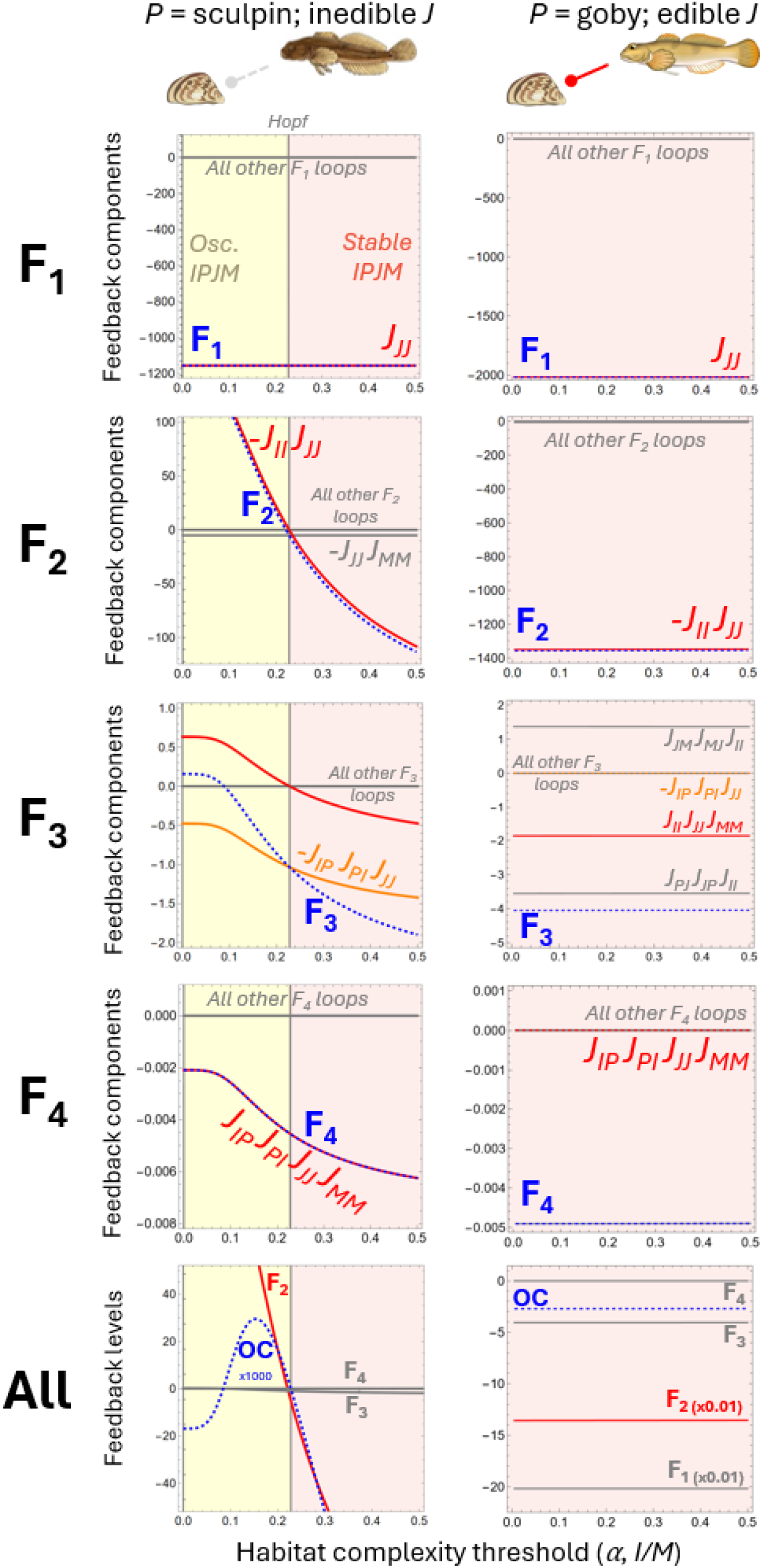
All feedback loops in the predator-prey and habitat complexity models according to habitat complexity threshold *α*. Colors and organization follow the conventions in Figure S2. This figure corresponds directly to Figure S3C & D and Figure S4. In all cases, *d*_*P*_ = 0.004, *k*_*D*_ = 200, and *ω* = 4 unless varied.

**Figure S6:**
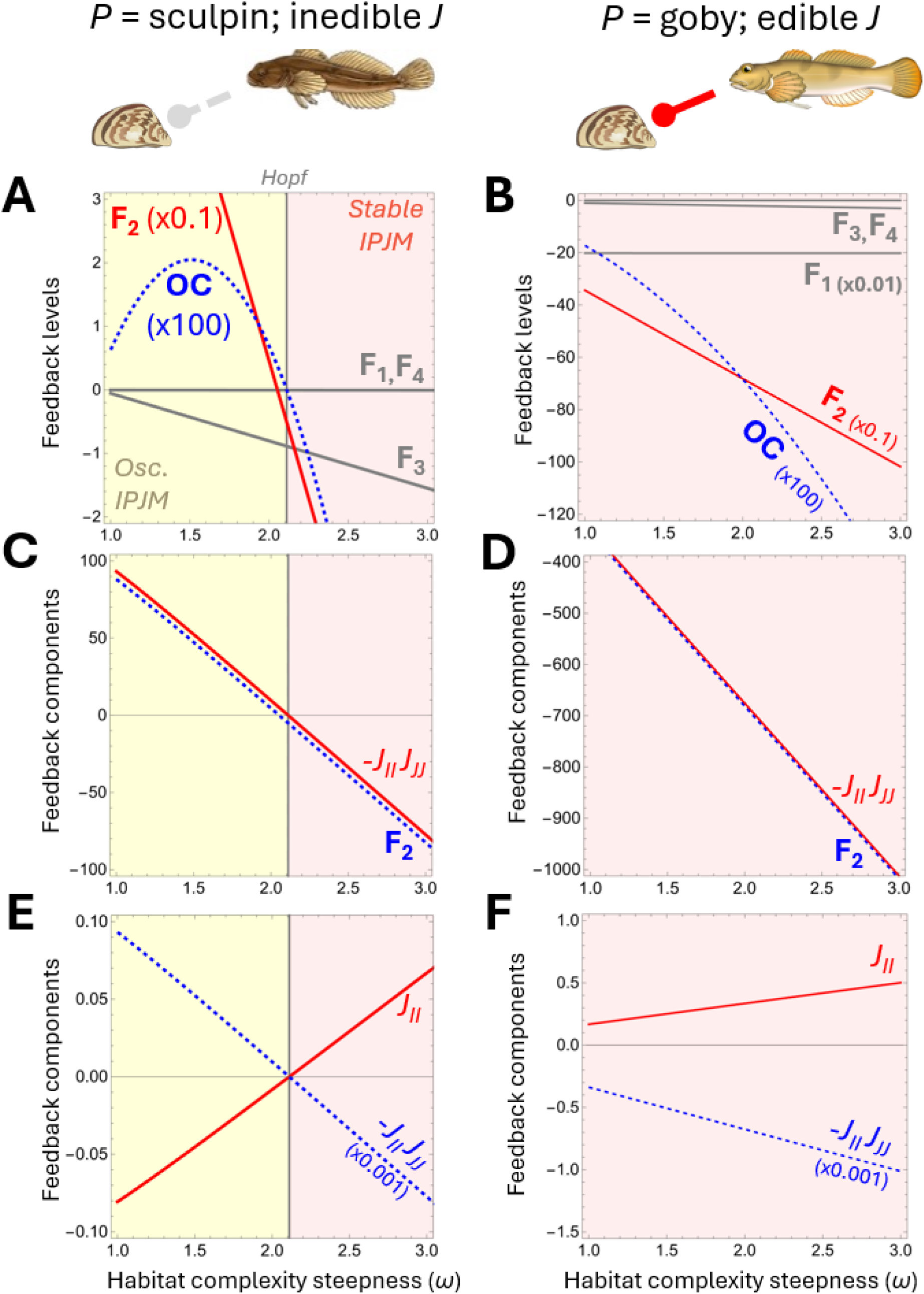
Loop tracing the onset and elimination of oscillations in *IPJM* models with sculpin and goby as the mid-trophic predator according to habitat complexity (HC) threshold steepness *ω*. Conventions follow Figures 4 and S4. **(A)** Without edible habitat complexity (EHC), the food web oscillates according to the criterion *OC* close to when **F**_**2**_ is positive. **(B)** With EHC, **F**_**2**_ still strengthens (becomes more negative) with increasing *ω*, but neither **OC** or **F**_**2**_ shifts or changes sign as with inedible *Dreissena*. **(C)** Without EHC, the self-effect of invertebrates *J*_*II*_ scaled by the self-effect of juvenile *Dreissena J*_*JJ*_ drives the behavior of **F**_**2**_, thus driving oscillations in the invertebrate-predator system. **(D)** With EHC, while *J*_*II*_ scaled by *J*_*JJ*_ does strengthen with *ω*, it is extremely stabilizing at all examined values. **(E)** Without EHC, *J*_*II*_ specifically undergoes the sign change relevant to food web stability, thus positive *J*_*II*_ drives oscillations with inedible *Dreissena*. **(F)** With EHC, *J*_*II*_ is positive and therefore destabilizing but is outweighed by other loops in the food web. In all cases, *d*_*P*_ = 0.004, *k*_*D*_ = 200, *α* = 1, and *ω* = 4 unless varied.

**Figure S7:**
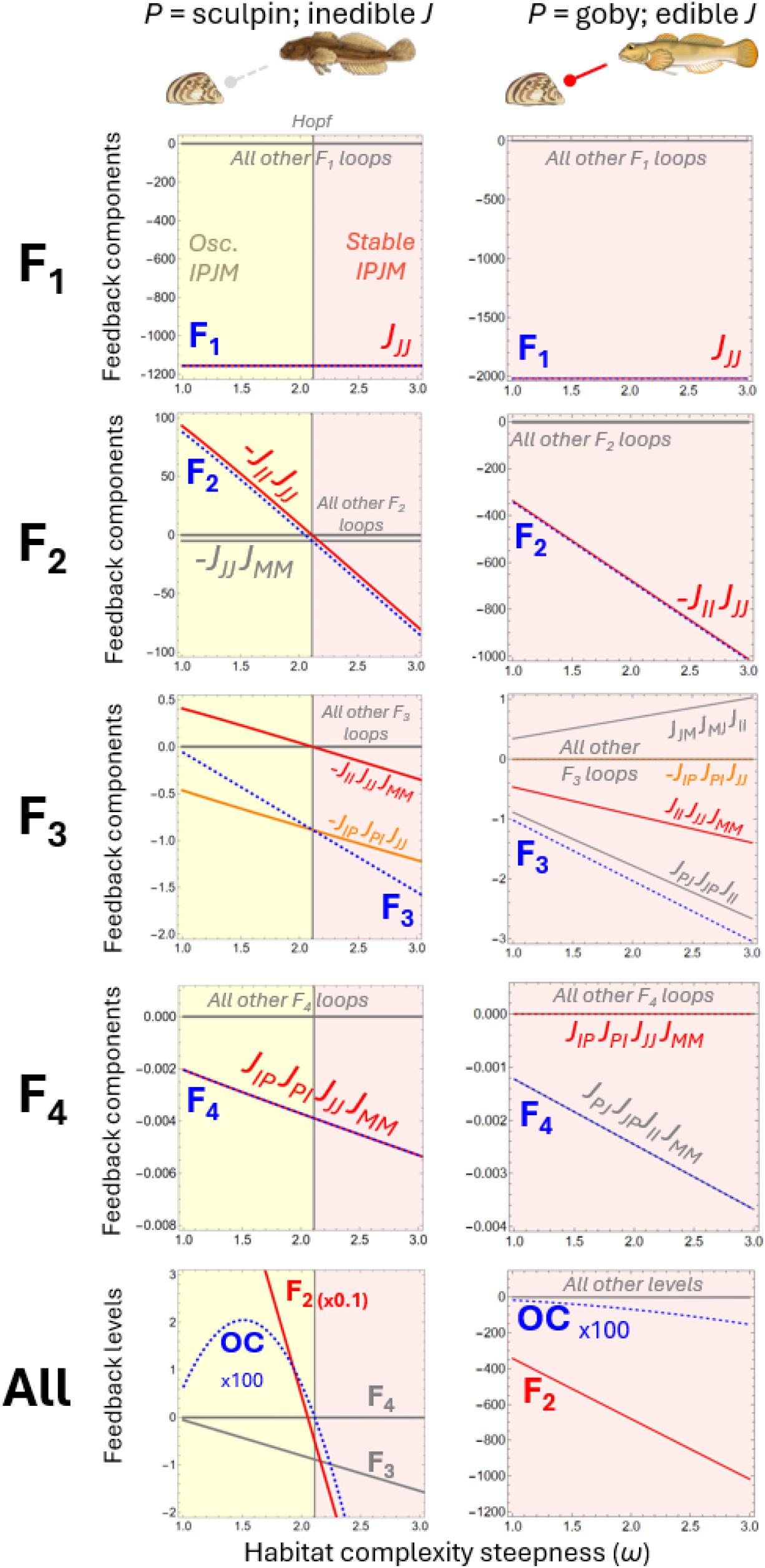
All feedback loops in the predator-prey and habitat complexity models according to habitat complexity steepness *ω*. Colors and organization follow the conventions in Figure S2. This figure corresponds directly to Figure S3E & F and Figure S6. In all cases, *d*_*P*_ = 0.004, *k*_*D*_ = 200, and *α* = 0.15. In all cases, *d*_*P*_ = 0.004, *k*_*D*_ = 200, *α* = 1, and *ω* = 4 unless varied.

**Figure S8:**
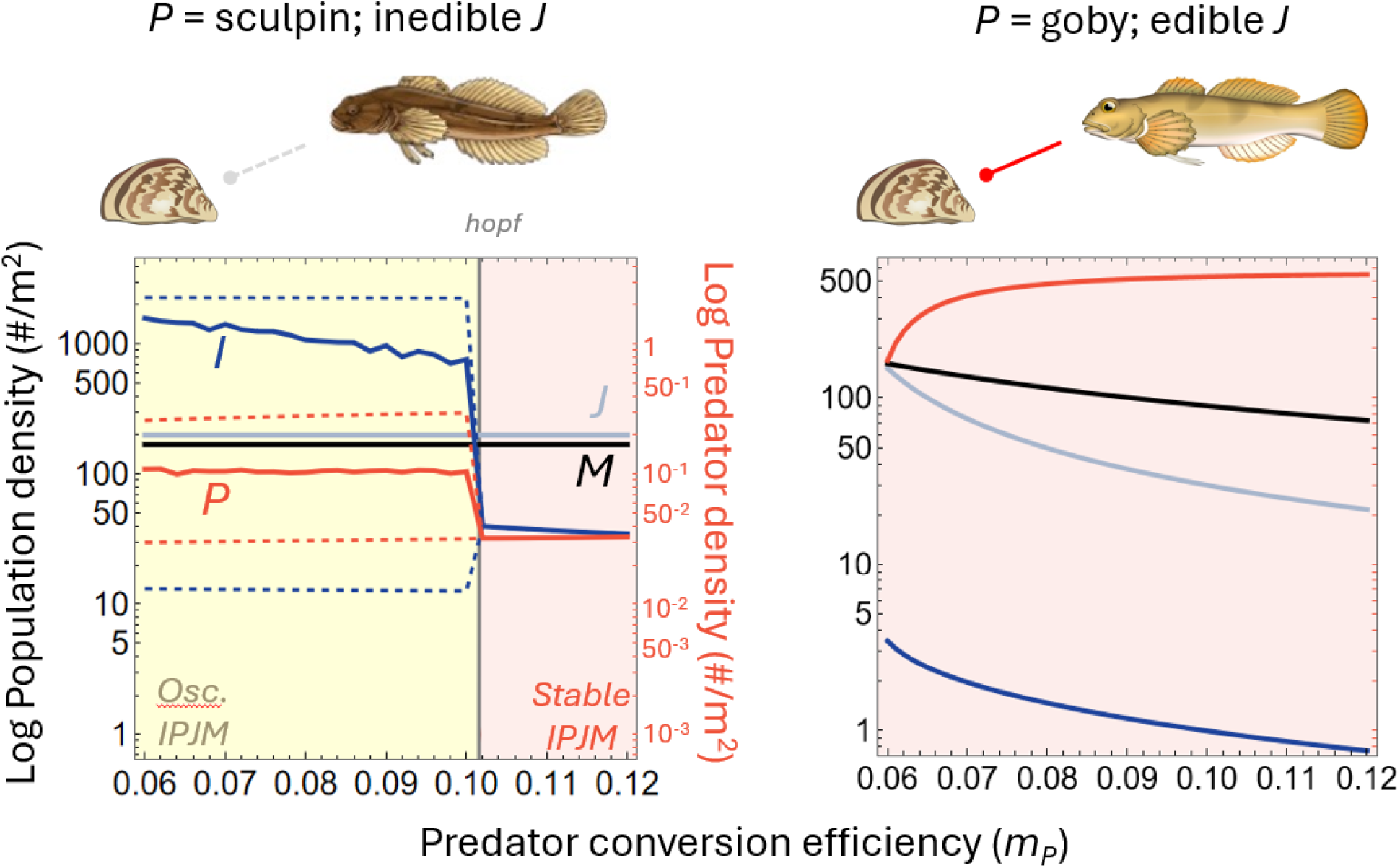
Population densities in the habitat complexity (HC) and edible habitat complexity (EHC) models according to changing predator conversion efficiency (*m*_*P*_). As in Figures 3 and 5 of the main text, in the oscillating region, dashed lines indicate the maxima and minima of the oscillations, while the solid line indicates the mean. Low conversion efficiencies can destabilize the food web in the HC model, but not the EHC model. In both cases, *d*_*P*_ = 0.004, *k*_*D*_ = 200, *α* = 0.15, and *ω* = 4.

